# Therapeutic Potential of a Tectorigenin Derivative (TED) in Herpetic Stromal Keratitis: Antiviral and lmmunomodulatory Mechanisms

**DOI:** 10.1101/2025.09.21.677568

**Authors:** Kailin Niu, Yang Hong, Mingming Yuan, Hong Zhang, Huixin Meng, Wei Yang, Chongjun Yuan, Yuan Wang, Hongling Xu, Jing Zhou, Lei Zhang

**Author notes:** Corresponding author: Jing Zhou, Lei Zhang, Tel.

## Abstract

**Objectives:** In this study, the anti-HSV-1 effects of TED were investigated *in vitro* and *in vivo*, its therapeutic efficacy against herpetic stromal keratitis (HSK) was evaluated, and its mechanism of action was examined.

**Methods:** We assessed the cytotoxicity and anti-HSV-1 activity of TED in Vero cells using CCK-8 and plaque assays. The HSK model mice received TED (2.5, 5, or 15 mg/kg), and symptom progression was monitored. The therapeutic effects were evaluated through HE staining, corneal immunofluorescence, and HSV-1 gB quantification. Mechanistic studies were performed to examine the expression of TLR pathway genes, CD4^+^/CD8^+^ T-cell infiltration in the cornea and draining lymph nodes, and dendritic cell activation.

**Results:** TED showed potent anti-HSV-1 activity *in vitro* and ameliorated HSK in mice, improving body weight and reducing corneal pathology and the incidence of encephalitis. It decreased HSV-1 loads in the cornea, trigeminal ganglion, and brain while modulating cytokines (downregulating IFN-α/γ, TNF-α, and IL-1β and upregulating IL-10). TED suppressed the TLR pathway (TLR2/3/9, TRAF6, IRF3, and NF-κB) in the corneal epithelium and reduced CD4^+^/CD8^+^ T-cell and dendritic cell infiltration while increasing the number of CD4^+^ T cells in the lymph nodes.

**Conclusions:** TED has significant anti-HSV-1 activity and effectively treats HSK by reducing viral loads, improving symptoms, and modulating immune responses through TLR pathway inhibition and immune cell regulation, suggesting its potential as an HSK therapeutic.

## Introduction

Herpes simplex virus keratitis (HSK), caused by herpes simplex virus-1 (HSV-1), is recognised as a leading cause of blindness worldwide. The current treatment options for HSK are largely limited to guanosine analogues, corticosteroids, and immunosuppressants^[1]^. However, these treatments are often associated with drug resistance and a high risk of disease recurrence. Hence, there is an urgent need to develop novel HSK therapeutics with enhanced efficacy and reduced adverse effects.

Natural medicinal compounds, particularly bioactive phytochemicals derived from plant sources, have demonstrated remarkable potential as antiviral therapeutics. Tectorigenin, a naturally occurring isoflavone primarily derived from the rhizomes of *Belamcanda chinensis* (L.) DC. (also known as Belamcandae Rhizoma), is a plant widely used in traditional Chinese medicine^[2]^. This compound exhibits a range of biological activities, including anti-inflammatory, antioxidant, and antimicrobial effects^[3–5]^. However, research on its antiviral properties remains limited. Among the few studies available, Jeong et al. (2015) demonstrated that tectorigenin could eliminate HIV-1-infected macrophages *in vitro*^[6]^. In addition to its antiviral potential, tectorigenin has shown anti-inflammatory effects in both *in vitro* and *in vivo* in models of *Dabie bandavirus* infection^[7]^. Additionally, Ponce et al.’s (2024) molecular docking simulations revealed that tectorigenin interacts with herpes simplex virus glycoprotein D—a key viral protein essential for HSV host cell entry^[8]^. Thus, tectorigenin might be a promising candidate for the development of therapeutic agents against HSV-1 infection.

To fully exploit the therapeutic potential of natural compounds, structural optimization through chemical modification has become an important strategy. This approach allows the enhancement of key pharmaceutical properties, including bioavailability, target specificity, and therapeutic potency, while maintaining the beneficial characteristics of the parent molecules^[4, 5]^. Studies have demonstrated that chemical modification of oleanolic acid from *Ligustrum lucidum* significantly enhances its activity against hepatitis B virus (HBV), including the inhibition of HBsAg and HBeAg secretion, as well as HBV DNA replication^[3]^. Similarly, juglanin, a glycosylated derivative of kaempferol obtained through acylation glycosylation, has stronger anti-SARS-CoV-1 activity than its parent compound kaempferol^[9]^. In our previous work, we developed a novel chemically modified derivative of tectorigenin, designated MJJ, which was specifically designed to improve its antiviral activity^[10]^. MJJ, protected by an international patent, has demonstrated potent anti-Coxsackievirus effects and has been included in the Pharmacopoeia of the People’s Republic of China as a therapeutic agent for diseases caused by Coxsackievirus, such as myocarditis, paediatric pneumonia, and hand-foot-and-mouth disease^[11]^. Building on this success, we synthesised a new tectorigenin derivative, designated TED, through further structural optimization. Given the broad-spectrum antiviral potential of many bioactive compounds^[12–14]^, the present study was designed to evaluate the antiviral effects of TED against HSV-1 and its therapeutic potential in HSK.

HSK is considered an immunopathological disorder primarily driven by excessive cytokine production from both innate and adaptive immune responses^[12]^—a process that may be pharmacologically targeted by TED. During HSV-1 infection, the innate immune response is rapidly activated through viral recognition by TLR2/3/9, triggering the TRAF6/IRF3-NF-κB pathway to produce inflammatory factors and type I interferons (IFNs). Given the central role of the TLR-TRAF6/IRF3-NF-κB axis in this process, it represents a potential intervention target for tectorigenin derivatives, as tectorigenin has been shown to reduce cytokine secretion by modulating the TLR/NF-κB pathways in mice and rats across various disease models^[7, 15]^. As a consequence of innate immune activation, HSV-1-infected dendritic cells (DCs) undergo maturation and migrate to draining lymph nodes (DLNs), where they further amplify proinflammatory cytokine and IFN production while initiating antigen-specific adaptive immunity. Adaptive immunity exacerbates HSK severity through DLN-derived CD4^+^ and CD8^+^ T cells that migrate to and infiltrate the cornea, where their local upregulation of inflammatory cytokines (e.g., IFN-γ and IL-17) and IFNs amplifies tissue damage. To date, studies on the effects of tectorigenin and its derivatives on DCs and CD4^+^ and CD8^+^ T cells are limited. Wang et al. (2020) reported that tectorigenin could alleviate inflammation by decreasing the aggregation of lymphocytes in an allergic asthma model of ovalbumin-sensitised guinea pigs^[16]^. However, Sajad et al. (2017) demonstrated that tectorigenin expands splenic CD4^+^ and CD8^+^ T-cell populations in healthy mice, resulting in elevated cytokine concentrations^[17]^. These conflicting reports suggest that the regulatory effect of tectorigenin on lymphocytes may depend on the baseline immunological state of the animals. Therefore, it is scientifically valuable to examine how tectorigenin and its derivatives modulate lymphocyte responses in the context of HSV-1 infection.

In this study, we implemented an integrated experimental strategy encompassing (1) *in vitro* assessment of direct antiviral activity through plaque reduction and viral replication assays, (2) *in vivo* validation of the therapeutic efficacy of TED using a murine HSK model, and (3) mechanistic dissection of TLR pathway modulation (TLR2/3/9-TRAF6/IRF3-NF-κB) and immune rebalancing (DC/T-cell dynamics and cytokine networks). The aims of these investigations were to establish TED as a therapeutic agent with both virological and immunoregulatory properties while providing evidence-based guidance for its clinical development in HSK treatment regimens.

## 1 Materials and Methods

### 1.1 TED

TED, which exhibited favourable water solubility, is a structurally modified derivative based on the tectorigenin scaffold. The specific synthetic route is withheld because it is pending patent protection.

### 1.2 Antiviral Activity *in vitro*

#### 1.2.1 Cell and Virus

Vero cells (GNO10) were obtained from the Cell Bank of the National Collection of Authenticated Cell Cultures (Shanghai, China). Vero cells were maintained in minimum essential medium (MEM, Jiangsu KGI Biotechnology Co., LTD, Jiangsu, China) containing 10% fetal bovine serum (gibco) at 37 ℃ with 5% CO_2_. The HSV-1 McIntyre strain (VR-539) was obtained from the American Type Culture Collection and propagated in Vero cells. The infectivity of the viral stock was determined by titration using two methods.: the 50% tissue culture infectious dose (TCID_50_) was determined using the Reed-Muench method^[18]^, while the plaque-forming unit (PFU) titer was determined by plaque assay under an agarose overlay, as described by Zhou et al. (2024)^[19]^.

#### 1.2.1 Cytotoxic Effect of TED

A CCK-8 assay was conducted to assess the cytotoxicity of TED and to determine its maximum non-toxic concentration prior to antiviral evaluation. In brief, Vero cells were seeded into 96-well plates at a density of 3×10^4^ cells per well in a final volume of 100 μL and incubated at 37 ℃ under 5% CO_2_. After 24 hours of incubation, the medium in each well was replaced with 300 μL of fresh medium containing TED at varying concentrations (0, 20, 40, 80, 160, or 320 μg/mL). After a further 72-hour incubation under the same conditions, the supernatant was removed, and a diluted CCK-8 solution (Dojindo, Kumamoto, Japan) was added to each well and incubated for another 1.5 h. Then, the plates were subjects to absorbance measurement at 450 nm with a microplate reader (Molecular Devices, California, USA).

#### 1.2.2 HSV-1 Infection *in vitro* and TED Treatment

To study the antiviral potential of TED, Vero cells were plated in 96-well plates according to the aforementioned protocol. Briefly, cells were seeded at 3×10⁴cells per well in 100 μL of culture medium and incubated overnight. Subsequently, the cells were inoculated with HSV-1 at a concentration of 100 TCID_50,_ 100μL/well for 1 h^[20]^ . After infection, the culture medium was replaced with 300 μL of one of the following: MEM (virus control), MEM containing 26 μg/mL acyclovir (ACV), or MEM containing TED at concentrations of 20, 40, 80, 160, and 320 μg/mL. Meanwhile, a separate uninfected group cultured in MEM served as the negative control. After a further 72 h incubation, viral-induced cytopathic effect was assessed using a CCK-8 assay to evaluate the protective efficacy of TED.

To further evaluate the antiviral efficacy of TED, a plaque reduction assay was conducted. Vero cells were seeded in 12-well plates at a density of 3.0×10⁵cells per well in 2 mL of MEM and cultured until confluent monolayers formed. All wells, except those assigned to the normal control group, were inoculated with 50 PFU of HSV-1 in 1 mL per well ^[21]^. Control wells received 1 mL of MEM without virus. After 1 h of adsorption at 37 ℃ under 5% CO_2_, the inoculum was aspirated. The cells were then overlaid with 2 mL per well of a 0.8% agarose mixture, prepared by combining equal volumes of liquefied 1.6% agarose and 2× MEM maintenance medium containing various concentrations of TED (0, 20, 40, 80 μg/mL) or 26 μg/mL ACV. Following solidification at room temperature, the plates were incubated for 72 h at 37 ℃with 5% CO_2_. Then, cells were fixed with ice-cold 4% paraformaldehyde at 4 ℃ for 12 h. The agarose overlay was carefully removed, and the cell monolayers were washed twice with DPBS. Cells were stained with 500 μL of 0.1% crystal violet solution per well for 15 min at room temperature. Finally, the plates were rinsed thoroughly with distilled water, air-dried, and viral plaques were enumerated. The inhibition percentage was calculated using the following formula: viral inhibition (%) = [1-(number of plaques) _inhibitor_/(number of plaques)_controls_] ×100^[22]^.

### 1.3 Antiviral Activity in the Mouse Model

All animal experiments were approved by the Animal Research Ethics Committee of the Sichuan Academy of Traditional Chinese Medicine and conducted under the guidance of the Experimental Animal Management Committee of the Sichuan Academy of Traditional Chinese Medicine and following the relevant regulations established by the Society for Vision and Ophthalmology Research (approval number: DWSYLL-2025-013).

#### 1.3.1 Animal Source and Feeding Conditions

Female BALB/c mice (6 weeks old, 18–22 g) with normal ocular development were purchased from Beijing Huafukang Biotechnology Co. Ltd. and maintained under specific pathogen-free (SPF) conditions at the Animal Experiment Center of the Sichuan Academy of Chinese Medicine (Chengdu, China) prior to the study.

#### 1.3.2 Establishment of the HSK Model and Drug Delivery

Following one week of acclimatization, fifty-four mice were randomly assigned to six experimental groups (n=9 per group): control (control mice given PBS via eye drops), HSK (HSK mice given PBS via eye drops), GCV (GCV mice given 1 mg/mL ganciclovir via eye drops), TED-L (mice given 2.5 mg/mL TED via eye drops), TED-M (mice given 5 mg/mL TED via eye drops), TED-H (mice given 15 mg/mL TED via eye drops). Anaesthesia was induced by the intraperitoneal administration of 0.4% sodium pentobarbital (40 mg/kg body weight). Except for the control group, all mice received viral inoculation, specifically, 1 μL of HSV-1 MacIntyre strain suspension (2×10³ PFU/mL) was injected into the corneal stroma using a sterile 10 μL microsyringe (Sangon Biotech, Shanghai, China) inserted through the conjunctival rim at a 45° angle to minimize tissue damage^[23]^. Twenty-four hours post infection, the mice were treated with either PBS, GCV or TED six times daily for nine consecutive days.

#### 1.3.3 Mouse Keratopathy Score

Ocular surface clinical signs^[24, 25]^: Eyes and periocular regions were directly observed and scored according to the following criteria:

**Table 1.** Grading criteria for HSV-1-induced corneal opacity.

| Score | Scoring criteria |
| --- | --- |
| 0 | no epithelial lesions or punctate lesions, no oedema or cloudiness of the stroma |
| 1 | mild swelling of the eyelids |
| 2 | moderate swelling with some crusting |
| 3 | eye swollen 50% shut with severe crusting |
| 4 | eye crusted shut |

**Table 2.** Grading criteria for HSV-1-induced corneal opacity.

| Score | Scoring criteria |
| --- | --- |
| 0 | no opacity |
| 1 | mild cloudiness with visible iris |
| 2 | moderate cloudiness with obscured iris |
| 3 | total corneal cloudiness with invisible iris |
| 4 | total opacity with no posterior view |

Corneal neovascularization: The cornea was divided into four equal quadrants. The length of neovessels in each quadrant was rated on a scale from 0–4. If the neovascularized area in a quadrant occupied up to 1/4 of that quadrant, it was given a score of 1 point^[24]^. The total neovascularization score for each cornea was calculated as the sum of the scores from all four quadrants.

### 1.4 H&E Staining of Mouse Corneal Tissues

On Day 10 postinfection, mouse eyeballs were harvested and fixed with an ocular fixative solution (Lilai Biotech Co., Ltd., Chengdu, China)^[27]^. The samples were then embedded in paraffin, sliced into 3-μm-thick sections, and deparaffinized with xylene (Jiangyuan Industrial Technology Co., Ltd., Wuxi, China). Next, the sections were stained with haematoxylin (Sigma‒Aldrich, St. Louis, MO, USA), differentiated in acid alcohol, and blued in a weak alkaline solution. Counterstaining was performed with alcoholic eosin (Bomei Biotech Co., Ltd., Hefei, China). Finally, the sections were mounted with neutral balsam (Sinopharm Chemical Reagent Co., Ltd., Shanghai, China). All the tissue sections were initially screened at low magnification, followed by detailed observation and image acquisition at 400× magnification to evaluate specific pathological changes.

### 1.5 Immunofluorescence Staining

Corneal whole-mount staining was carried out as previously described. The paraffin sections were deparaffinized, subjected to antigen heat retrieval, permeabilized with 3% hydrogen peroxide, and blocked with 3% bovine serum albumin (BSA; Servicebio, Wuhan, China). The sections were then incubated overnight with the following primary antibodies: anti-CD31 antibody (1:200 dilution; Abcam, Cambridge, MA, USA) and HSV-1 gD antibody (1:100 dilution; Santa Cruz Biotechnology, Dallas, TX, USA). The sections were subsequently incubated for 30 minutes with the following secondary antibodies: FITC-conjugated goat anti-mouse IgG (1:100 dilution; Servicebio, Wuhan, China) and CY3-conjugated goat anti-rabbit IgG (1:200 dilution; Servicebio, Wuhan, China). Nuclei were counterstained with DAPI (Servicebio, Wuhan, China), and the sections were mounted with antifade mounting medium to prevent fluorescence quenching.

### 1.6 Immunohistochemical Staining

The paraffin sections were deparaffinized, permeabilized, blocked, and then incubated overnight at 4 °C with the following primary antibodies: CD4 (1:400 dilution; Abcam, Cambridge, MA, USA), CD8 (1:400 dilution; Abcam, Cambridge, MA, USA), and CD11c (1:100 dilution; Huabio, Hangzhou, China). Following primary antibody incubation, the sections were incubated with an HRP-conjugated goat anti-rabbit secondary antibody (Servicebio, Wuhan, China). Colour development was performed via freshly prepared 3,3’-diaminobenzidine substrate (Zhongsui Jinqiao Biological Co., Ltd., Beijing, China) at room temperature. Staining observation and analysis were performed using a Digital Trinocular Camera Microscope (Motic, Fujian, China) and a Data Image Analysis System (Media Cybernetics, Maryland, USA).

### 1.7 Flow Cytometry

The draining lymph nodes (DLNs) were carefully rinsed in 60 mm glass Petri dishes containing 2 mL of ice-cold PBS (Pernoside, Wuhan, China). The tissues were minced using sterile ophthalmic scissors and gently homogenized with a tissue grinder. The resulting cell suspension was filtered through a 200 μm nylon mesh, and the cell pellet was collected by centrifugation at 300 × g for 5 minutes at 4 °C. The pellet was resuspended in 100 μL of PBS and stained with the following fluorescently labelled antibodies: FITC-conjugated anti-mouse CD4 antibody (0.5 μL per test; BioLegend, San Diego, CA) and PE-conjugated anti-mouse CD8 antibody (1.25 μL per test; BioLegend, San Diego, CA). Staining was performed at room temperature for 30 minutes in the dark. After incubation, the cells were washed three times with PBS and finally resuspended in 300 μL of PBS for flow cytometric analysis. Data analysis was performed using CytExpert.

### 1.8 Quantitative Real-Time Polymerase Chain Reaction(qRT-PCR)

Total RNA was extracted from mouse tissues using the Molpure^®^ Cell/Tissue Total RNA Kit (TaKaRa, Japan). cDNA was synthesised by reverse transcription of 1 µg of total RNA using the PrimeScript™ RT Reagent Kit (TaKaRa, Japan) according to the manufacturer’s instructions. The reaction was carried out in a TCA0096 thermal cycler (Thermo Fisher Scientific, USA) with an incubation step at 37 °C for 15 min and 85 °C for 5 s.

Quantitative PCR (qPCR) was performed using TB Green™ Premix Ex Taq™ II (Tli RNaseH Plus) to assess the expression levels of HSV-1 gB glycoprotein and multiple cytokines, including IFN-α, IFN-γ, IL-1β, IL-10, IRF3, NF-κB, TLR2, TLR3, TLR9, TNF-α, and TRAF6. The amplification protocol consisted of initial denaturation at 95 °C for 30 sec, followed by 45 cycles of denaturation (95 °C, 5 sec), annealing (55 °C, 30 sec), and extension (72 °C, 30 sec), with fluorescence acquisition at each cycle. Postamplification melt curve analysis was conducted to confirm PCR product specificity. The primer sequences for each target gene are listed below:

**Table 3.**
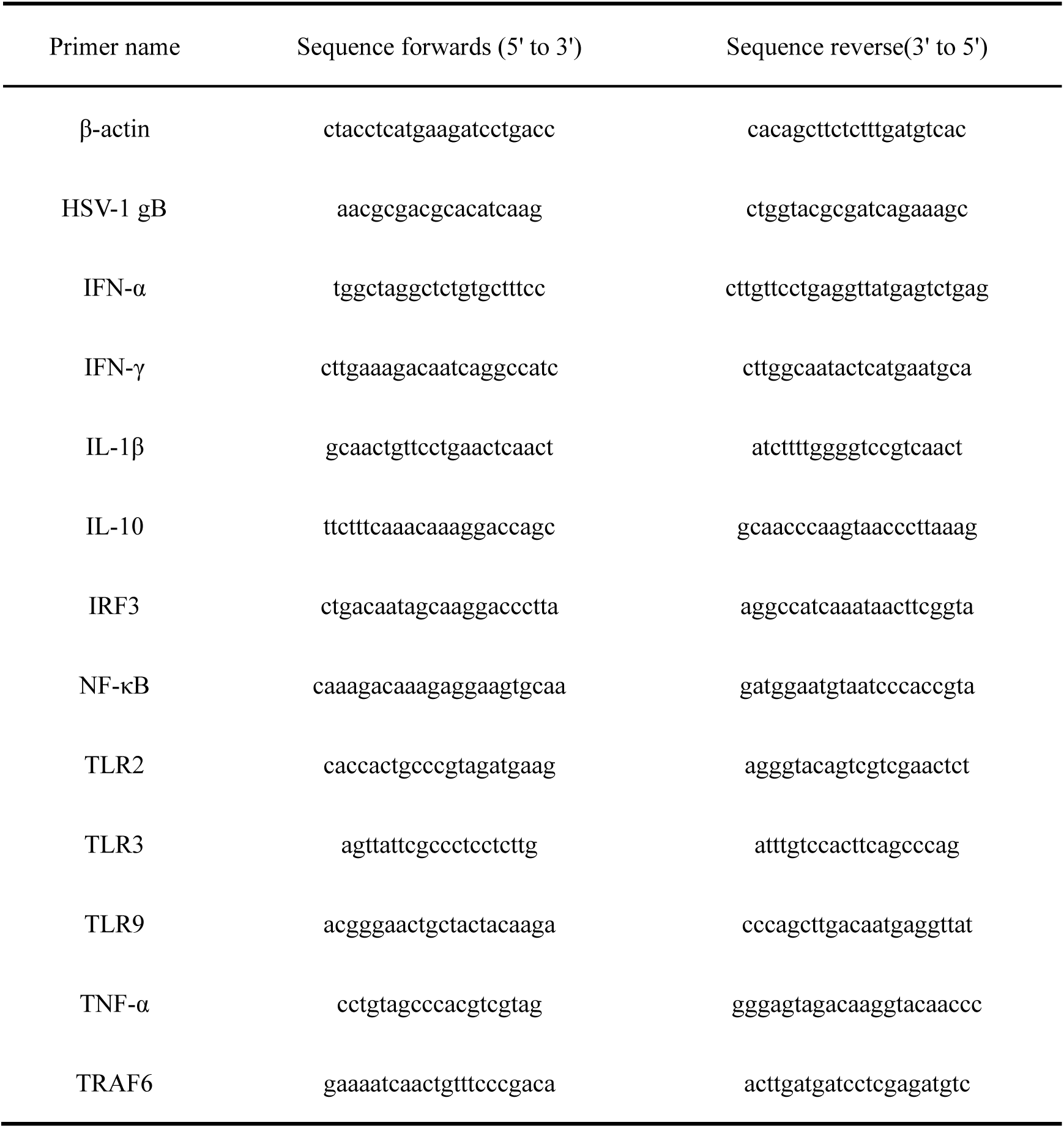
Primers used for qRT‒PCR.

### 1.9 Statistical Analysis

Statistical analysis was performed using GraphPad Prism 8.0 software. The data are presented as the means ± standard deviations (SDs) from at least three biological replicates. Student’s *t* test was used to compare differences between two groups, whereas one-way analysis of variance (ANOVA) was used for comparisons among three or more groups. Statistical significance was defined as follows: \**P* < 0.05, \*\**P* < 0.01, and \*\*\**P* < 0.001.

## 2 Results

### 2.1 TED Exhibits *In Vitro* Anti-HSV-1 Activity

The results after Vero cells were treated with various concentrations of TED for 72 h are shown in Fig. 1A. TED exhibited no significant cytotoxicity at concentrations below 320 μg/mL. Next, the inhibitory effect of TED on HSV-1 was evaluated. As shown in Fig. 1B, the results of the CCK-8 assay revealed that the OD value of the HSV-1-infected model group was significantly lower than that of the control group (*P* < 0.001), confirming successful viral infection in Vero cells. In the TED-treated groups, the inhibition of HSV-1-induced cytopathic effects was dose dependent, with an inhibition rate of 52% observed at 80 μg/mL. The results of the plaque assay further demonstrated the anti-HSV-1 activity of TED *in vitro* (Fig. 1B). Compared with the model group, TED inhibited HSV-1-induced vacuole formation in a dose-dependent manner, with the 80 μg/mL group showing a significant reduction (*P* < 0.05).

**Figure 1.**
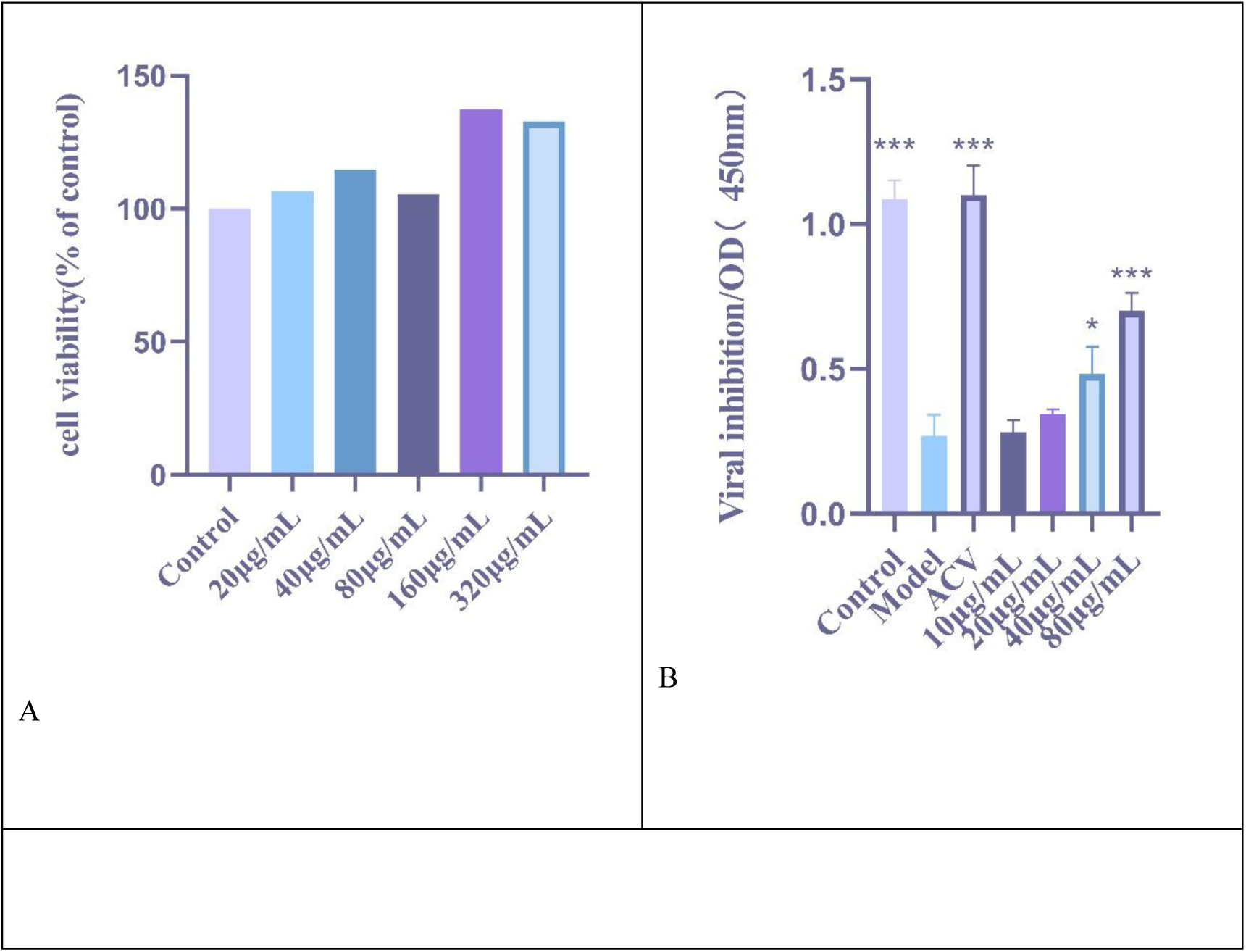
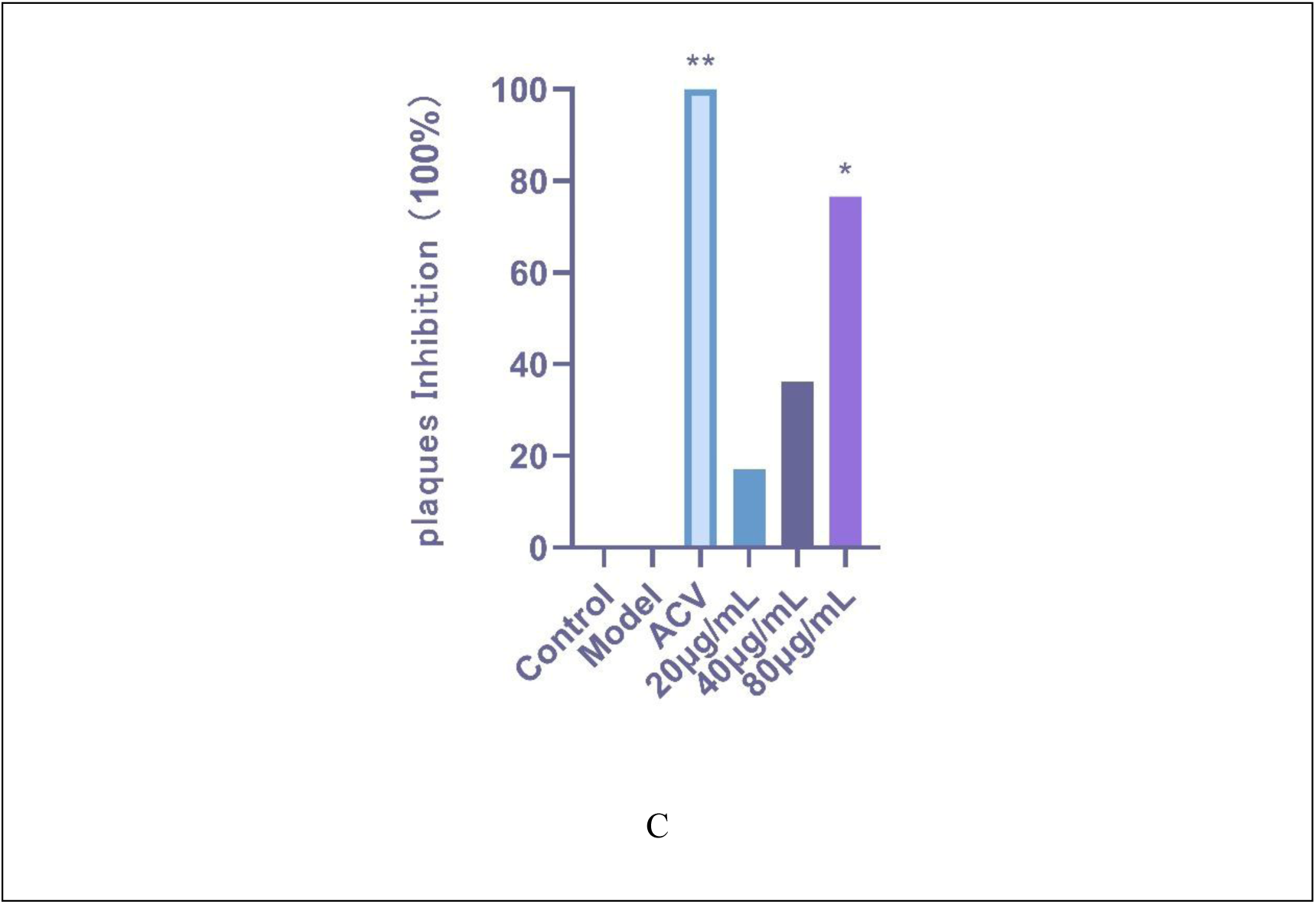
Cytotoxicity of TED in Vero cells and its inhibitory effect on HSV-1 infection. (A) Viability of Vero cells exposed to different concentrations of TED for 72 h. (B) Inhibitory effect of TED on HSV-1 infection in Vero cells (CCK-8 assay). (C) TED-mediated inhibition of plaque formation. Data are shown as the means ± SDs of three separate experiments. (\**P* < 0.05, \*\**P* < 0.01, \*\*\**P* < 0.001).

### 2.2 TED Inhibited HSK and Reduced the HSV-1 Viral Load in a Mouse Model

Compared with the control group, the HSK model group exhibited weight loss and a significant difference (*P* < 0.01), while TED administration suppressed this trend (Fig. 2B). The progression of clinical manifestations in the HSK group was as follows: on Day 2 postinfection, periocular secretions and mild eyelid swelling were observed; on Day 4, the model group mice exhibited pupil dilation, initial corneal opacity, and early neovascularization at the corneal‒conjunctival junction; and on Day 9, severe corneal opacity with obscured posterior view was evident, accompanied by extensive neovascularization extending from the corneal periphery towards the centre (Fig. 2A). Additionally, the model group of mice developed encephalitis symptoms, including emaciation, a hunched posture, head tilt, circling behaviour, hyperactivity, and a heightened startle response, indicating viral spread along the trigeminal pathway from the cornea to the central nervous system. In contrast, TED-treated mice presented significant improvements in clinical symptoms, with reduced corneal opacity, attenuated neovascularization, and an absence of neurological manifestations (Fig. 2A).

**Figure 2.**
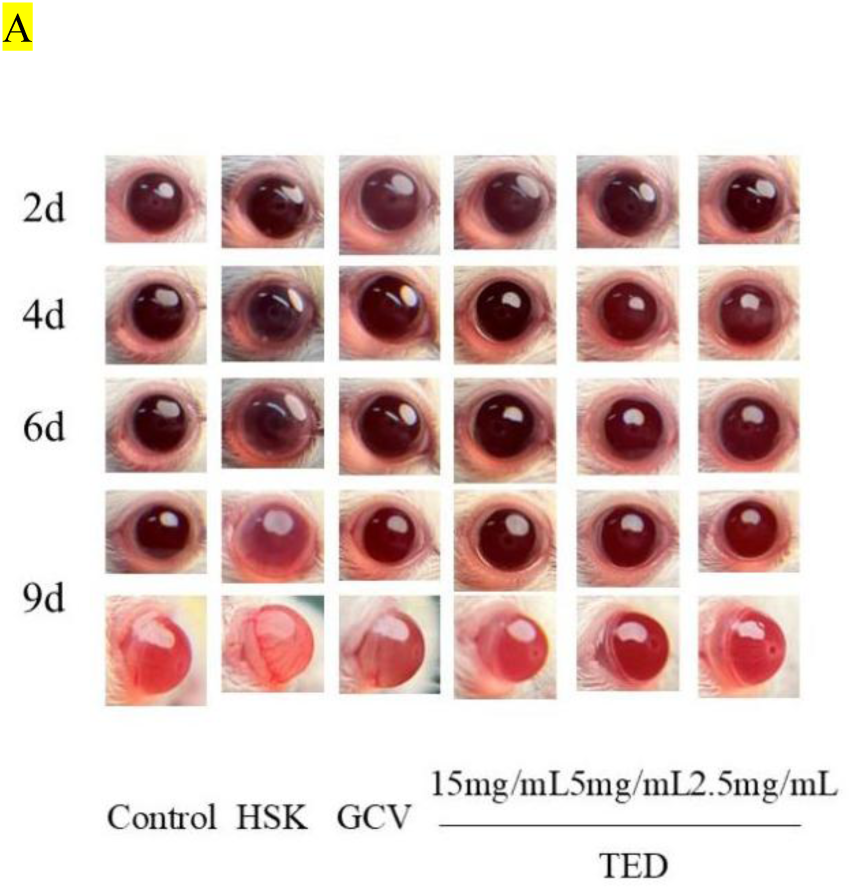
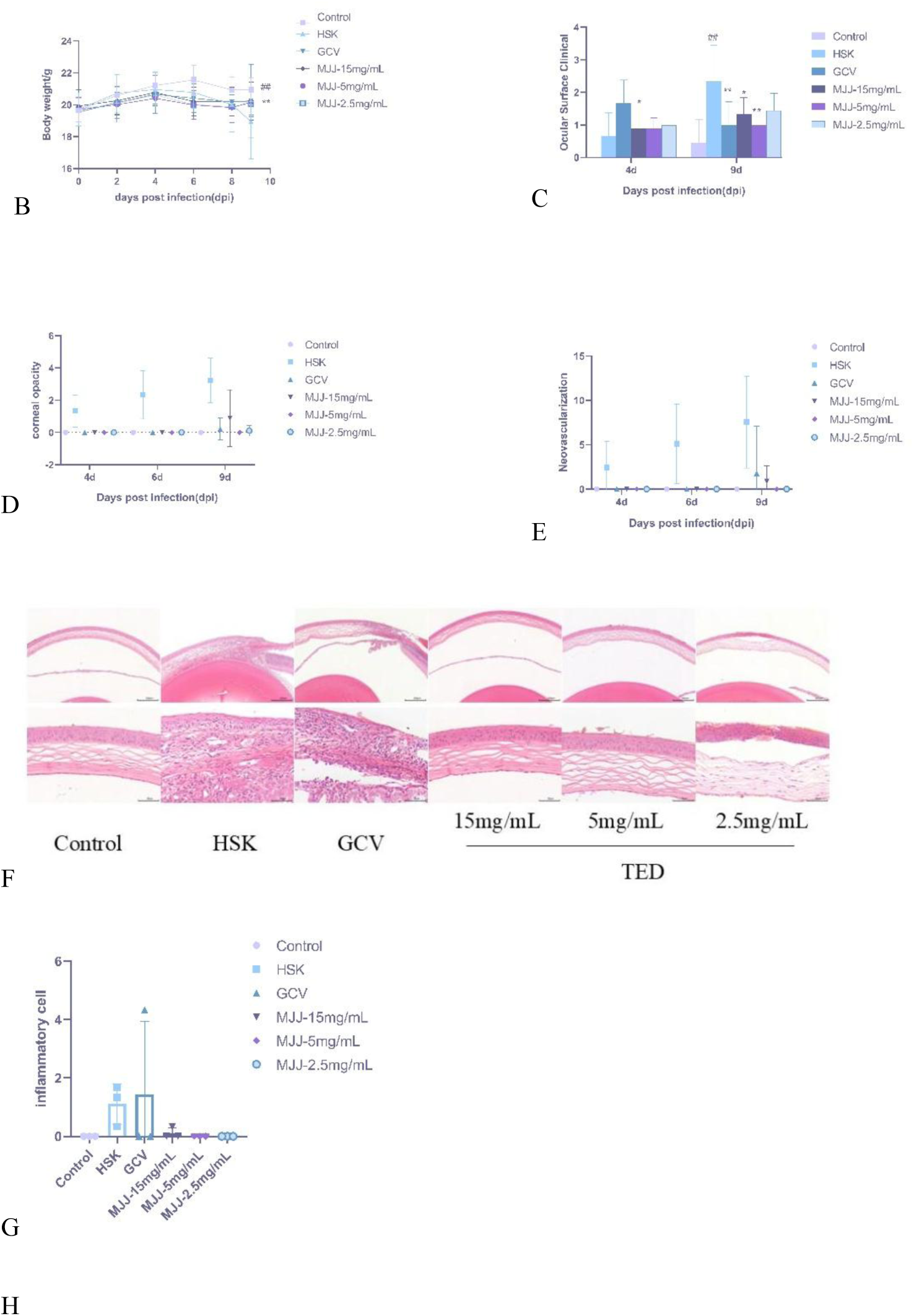
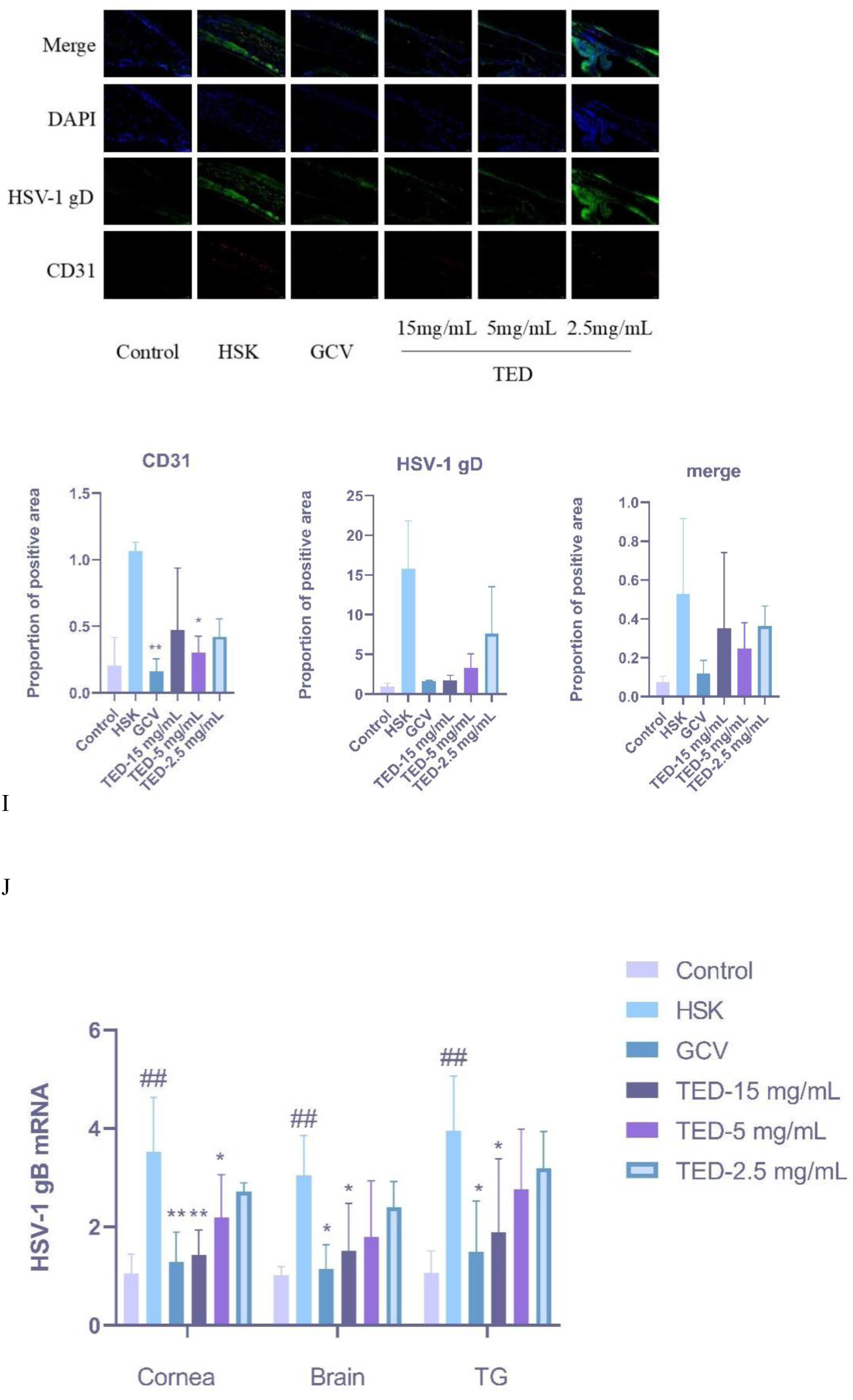
Inhibitory effects of TED on HSK progression and the HSV-1 viral load. (A) Ocular surface morphology in mice. (B) Body weight variation in mice (n=9). (C) Ocular surface clinical scoring results for the mice (n=9). (D) Corneal opacity scoring results for the mice (n=9). (E) Corneal neovascularization scoring results for the mice. (n=9). (F-G) HE staining of mouse corneal sections and quantification of leukocyte infiltration. (H) Representative micrographs of HSV-1 gD (green) alongside CD31 (red, a marker of neovascularization)-positive endothelial cells in the cornea (n=3). (I) The density of HSV-1 gD and CD31. (J) qRT‒PCR analysis of the HSV-1 viral load in the cornea, TG, and brain (n=3). The data are presented as the means ± SDs of three independent experiments. (\**P* < 0.05, \*\**P* < 0.01, \*\*\**P* < 0.001)

Quantitative assessment of clinical symptoms using ocular surface scoring, corneal opacity grading, and neovascularization area measurement revealed significant differences among the groups. On Day 9 post infection, compared with the control group, the model group presented significantly higher ocular surface clinical scores (*P* < 0.001). Treatment with Ganciclovir(GCV) and varying doses of TED resulted in a substantial reduction in ocular surface score (*P* < 0.05) (Fig. 2C). In the HSK model group, corneal opacity and neovascularization progressively worsened over time. In contrast, both the TED-5 mg/mL and TED-2.5 mg/mL groups presented no corneal turbidity or neovascularization throughout the entire observation period (Fig. 2D-E).

Histopathological evaluation of corneal tissues revealed distinct morphological differences across the experimental groups. The control group displayed intact corneal architecture with no significant pathological changes. In contrast, the HSK model group presented hallmark pathological features, including focal degeneration and necrosis of corneal epithelial cells, disorganization of stromal collagen fibres, and moderate inflammatory cell infiltration. Notably, rod-shaped neutrophils were present within the stroma, along with fibrotic tissue proliferation and early capillary formation. TED treatment conferred dose-dependent protective effects. All the TED-treated groups maintained a well-preserved corneal structure with only minor pathological alterations. The TED-15 mg/mL group presented mild inflammatory cell infiltration but no significant tissue degeneration or necrosis. Moreover, the TED-2.5 mg/mL and TED-5 mg/mL groups retained nearly normal corneal morphology, with no observable inflammatory infiltration or structural abnormalities (Fig. 2 F-G).

To further evaluate the antiangiogenic effects of TED in HSV-1-induced HSK, this study employed immunofluorescence staining to assess the expression of corneal HSV-1 gD and CD31 (a marker of corneal neovascularization). The results revealed increased expression of HSV-1 gD and CD31 in the HSK model group. In contrast, TED treatment led to a dose-dependent reduction in HSV-1 gD expression. Consistent with these findings, treatment with 5 mg/mL TED significantly reduced CD31-positive expression (*P* < 0.05) (Fig. 2H-I).

Viral load quantification in corneal, brain, and TG tissues on Day 10 revealed significantly higher HSV-1 gB levels in the model group than in the control group (*P* < 0.05). All the TED treatment groups demonstrated a dose-proportional reduction in the HSV-1 gB load across all the examined tissues (*P* < 0.05 or *P* < 0.01) (Fig. 2J).

### 2.2 TED Reduces Corneal Surface Inflammatory Cytokines

qRT‒PCR analysis revealed that HSV-1 infection significantly upregulated the expression of corneal inflammatory cytokines, including TNF-α, IFN-α, IFN-γ, and IL-1β (*P* < 0.01). Conversely, TED administration resulted in a reduction in these mediators (Fig. 3A–D). HSV-1 infection led to a significant reduction in the expression of IL-10 mRNA (*P* < 0.01), while TED-15 mg/mL effectively attenuated this inhibitory effect (*P* < 0.05) (Fig. 3E).

**Figure 3.**
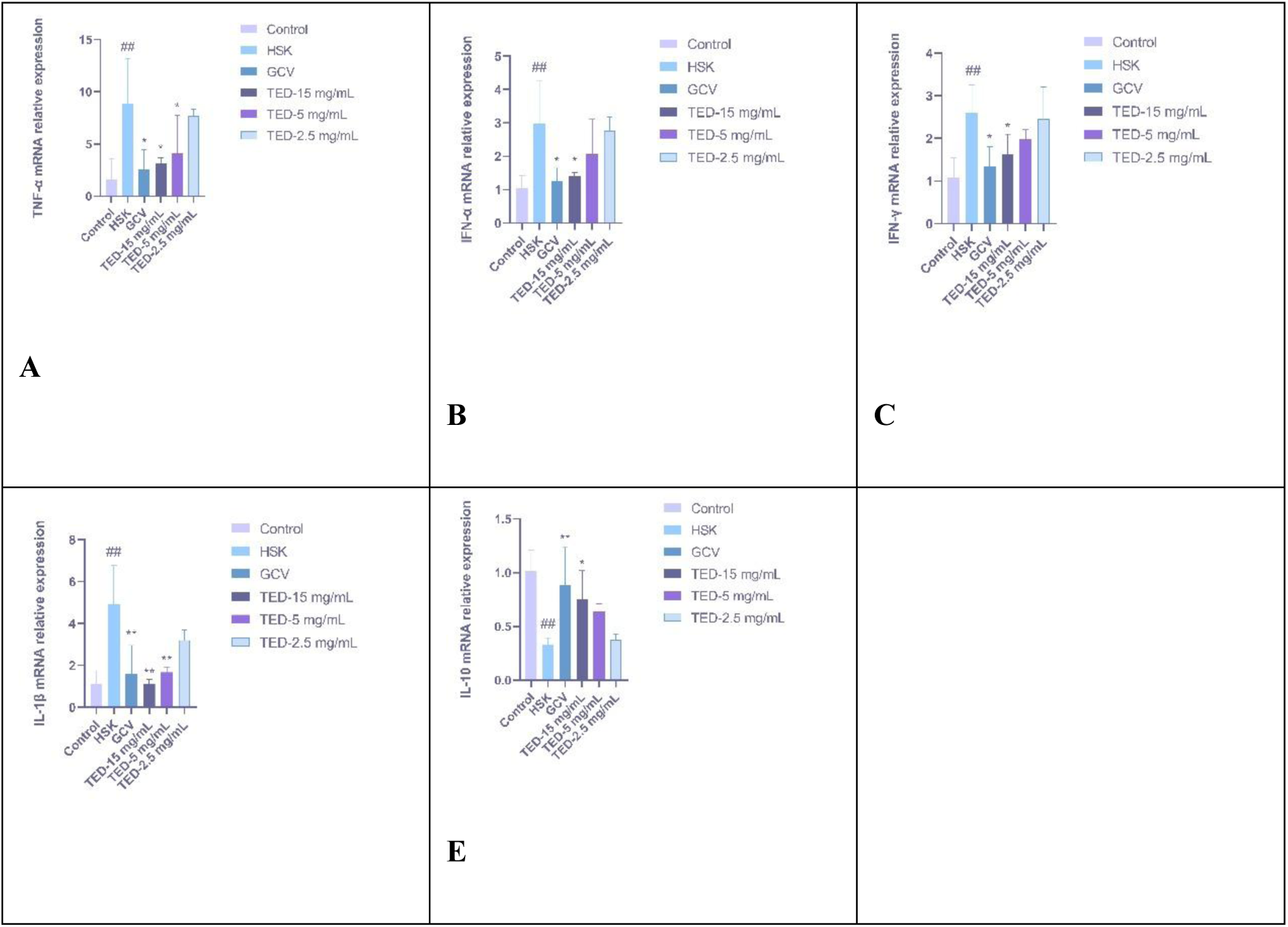
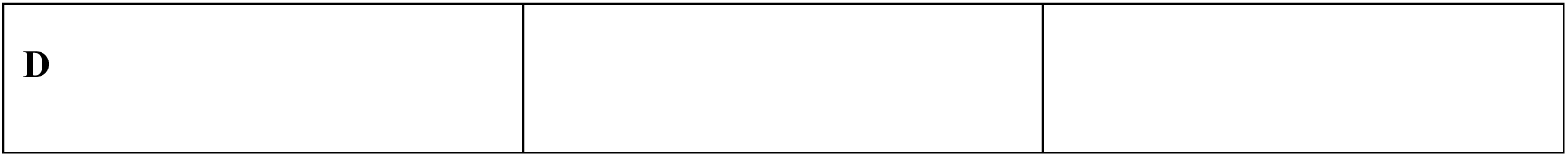
TED treatment significantly regulates proinflammatory cytokine mRNA expression levels on the corneal surface. (A-E) qRT‒PCR analysis of corneal TNF-α, IFN-α, IFN-γ, IL-1β, and IL-10 expression levels in mice (n = 3). The data are presented as the means ± SDs of three independent experiments (\**P* < 0.05, \*\**P* < 0.01, \*\*\**P* < 0.001).

### 2.3 TED Modulated Toll-like Receptor Signalling in the Mouse Cornea

The corneal tissues of HSK mice presented markedly elevated TLR2/3/9 pathway-related mRNA expression, with TLR2, TLR3, and TLR9 mRNA levels significantly exceeding those in healthy controls. (*P* < 0.01). Corresponding increases in gene expression were observed for downstream signalling molecules, including TRAF6, IRF3, and NF-κB (*P* < 0.01 or *P* < 0.05) (Fig. 4A-F). Compared with those in the HSK group, the levels of TLR2, TLR3, and IRF3 mRNA expression in the TED 5 mg/mL and 15 mg/mL groups were significantly lower (P < 0.01 or P < 0.05). Compared with the model group, the TED 15 mg/mL group presented inhibited TLR9, TRAF6, and NF-κB mRNA expression (*P* < 0.01 or *P* < 0.05).

**Figure 4.**
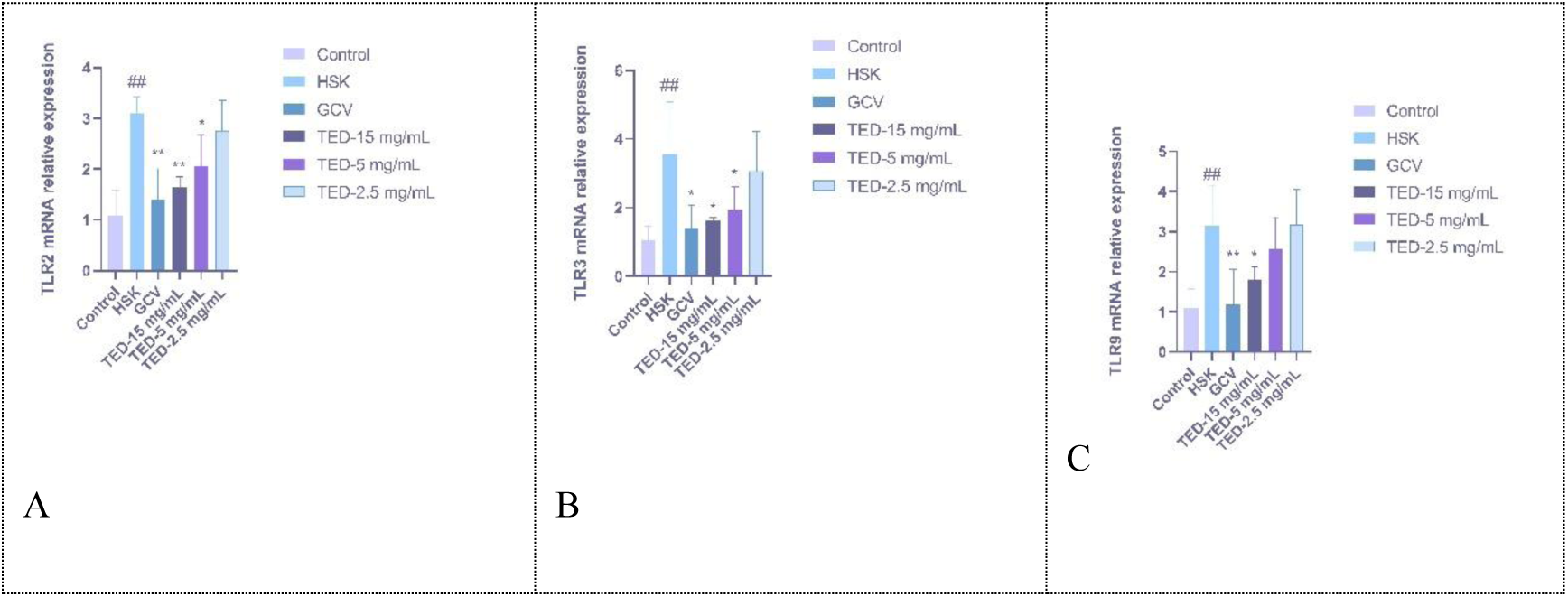
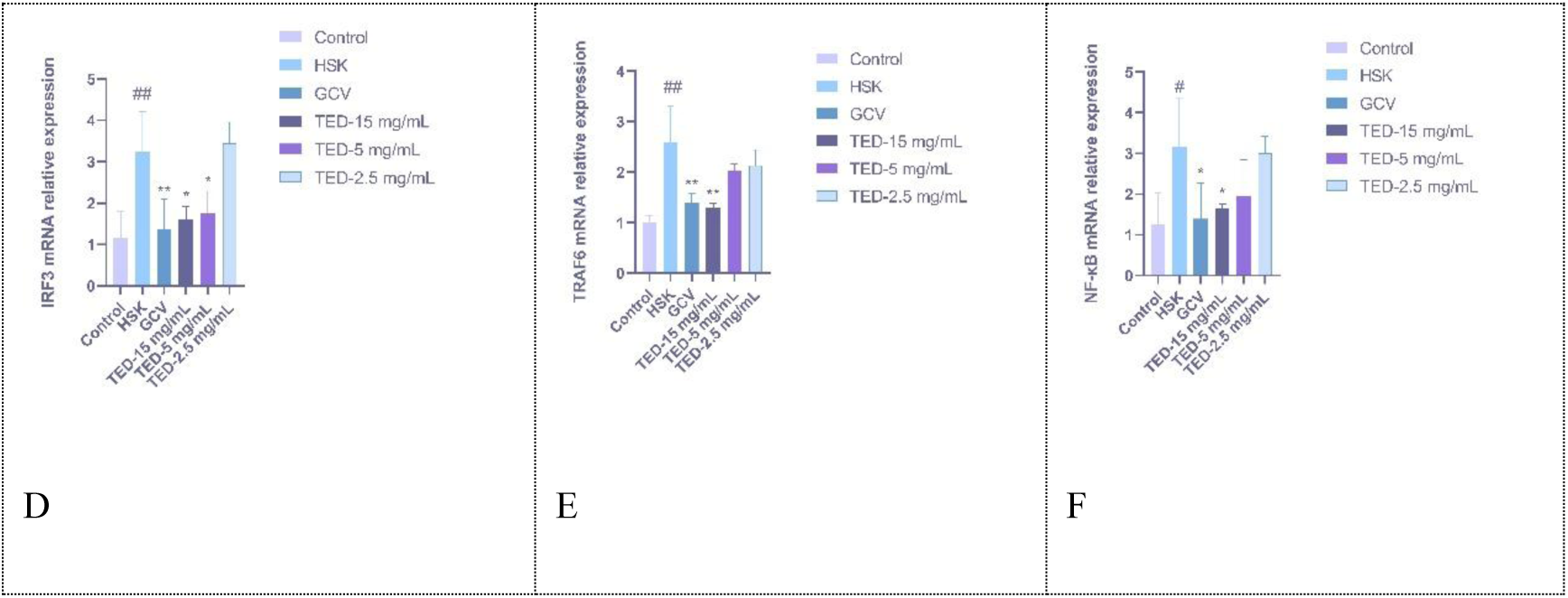
TED inhibits TLR2/3/9 pathway-related mRNA expression in the mouse cornea. (A-F) qRT‒PCR analysis of TLR2, TLR3, TLR9, TRAF6, IRF3, and NF-κB mRNA expression in corneal tissues (n = 3). The data are presented as the means ± SDs of three independent experiments. (\**P* < 0.05, \*\**P* < 0.01, \*\*\**P* < 0.001)

**Figure 5.**
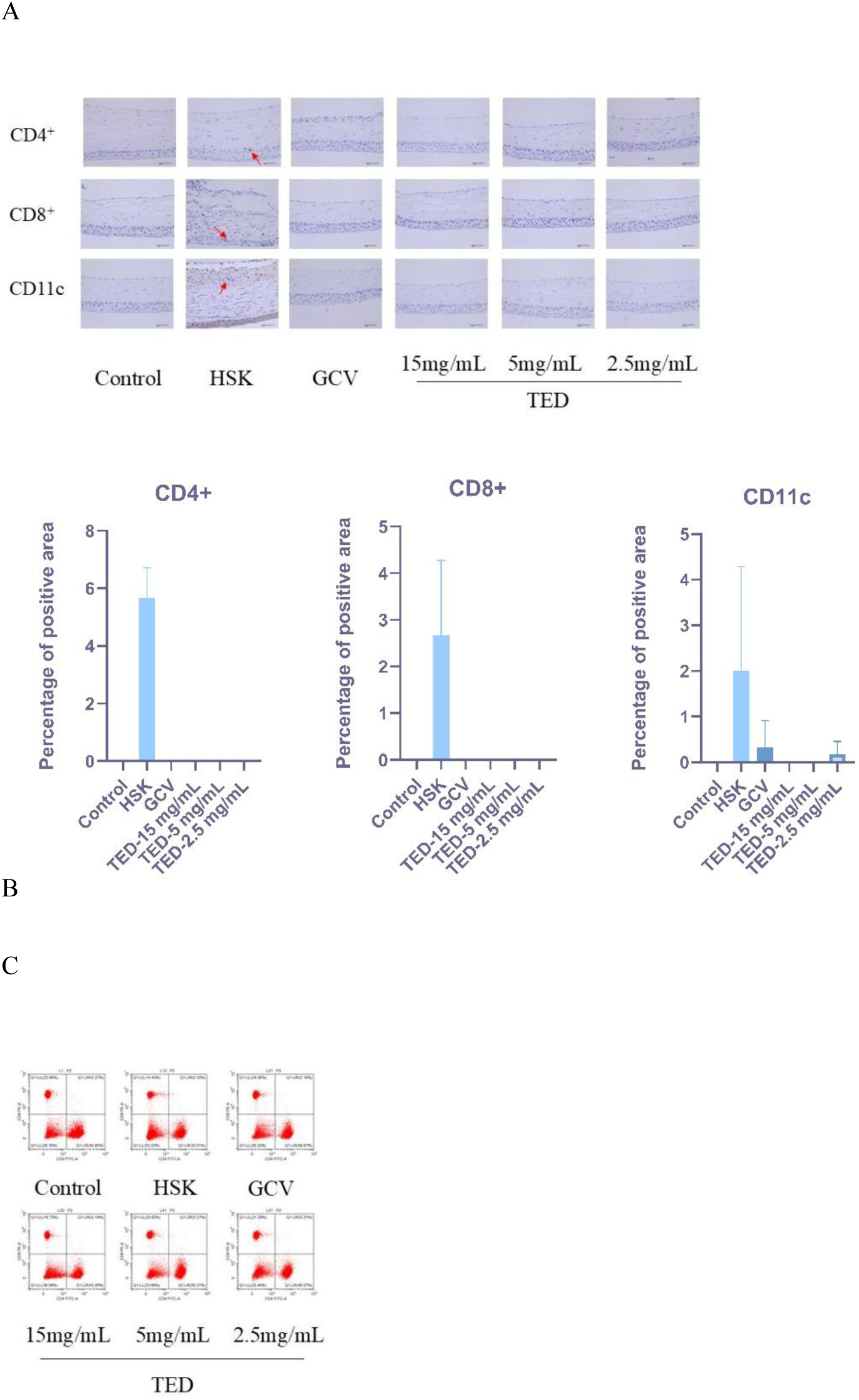
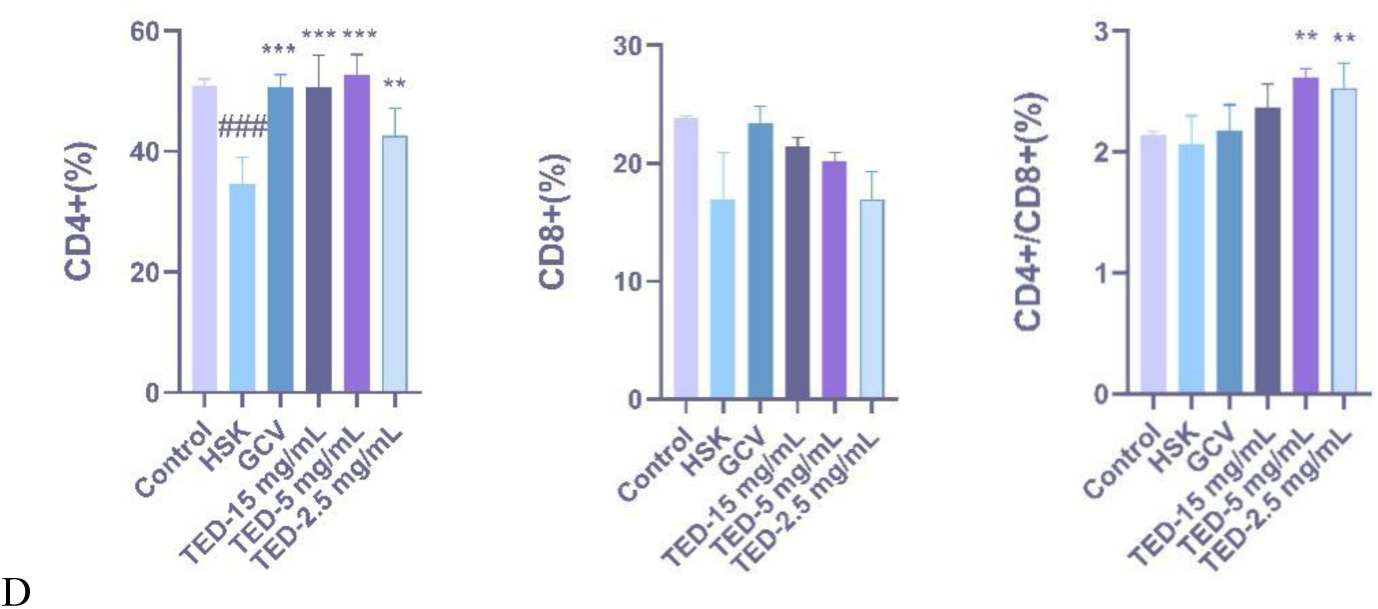
TED treatment regulates CD4+ and CD8+ T cells and dendritic cells during HSV-1 infection. (A-B) Immunofluorescence staining results revealed CD4^+^ and CD8^+^ T lymphocytes as well as CD11c^+^ dendritic cells in corneal tissues. Red arrows highlight positively stained cells exhibiting specific fluorescence signals for each marker. (C-D) Dynamic changes in CD4^+^ helper and CD8^+^ cytotoxic T-cell frequencies and the CD4^+^/CD8^+^ T-cell ratio within DLNs. The data are presented as the means ± SDs of three independent experiments (\**P* < 0.05, \*\**P* < 0.01, \*\*\**P* < 0.001).

### 2.5 TED Modulates Adaptive Immune Responses in HSV-1 Infection in the Mouse Cornea

We hypothesized that TED attenuates HSK development by selectively modulating CD4^+^ and CD8^+^ T-cell responses in the cornea and DLNs, thereby rebalancing adaptive immune mechanisms in corneal inflammation. Accordingly, we performed immunohistochemical analysis of CD4^+^ and CD8^+^ T cells and the DC marker CD11c in the cornea and flow cytometric analysis of immune cell populations, including CD4^+^ and CD8^+^ T lymphocytes, along with quantification of CD4^+^/CD8^+^ T-cell ratios in DLNs. Immunohistochemical analysis revealed significant infiltration of CD4^+^ and CD8^+^ T cells, along with the DC marker CD11c, in HSK corneas compared with normal control corneas. In contrast, the TED-treated groups showed a complete absence of T-cell infiltration, with the 5 mg/mL and 15 mg/mL doses preventing dendritic cell recruitment. Compared with that in the control group, a significant reduction in the CD4^+^ T-cell frequency was observed in the DLNs of the HSK group (*P* < 0.01). Conversely, TED administration resulted in an increase in CD4^+^ T-cell levels (*P* < 0.01).

## 3 Discussion

HSK remains a significant cause of infectious blindness worldwide, with current treatment options limited by drug resistance and frequent recurrence. While tectorigenin, a natural isoflavone, has demonstrated anti-inflammatory and antimicrobial properties, its potential antiviral effects against HSV-1 have not been fully characterized. Preliminary evidence suggests that tectorigenin may interact with HSV glycoprotein D. Building on our previous success with MJJ, a patented tectorigenin derivative clinically approved for Coxsackievirus infections, we hypothesized that structural optimization of tectorigenin could also yield effective anti-HSV-1 agents. In this study, we systematically evaluated the effects of TED on HSV-1 infection and HSK pathogenesis using both *in vitro* and murine models.

We initially investigated the antiviral activity of TED against HSV-1 in cell culture systems. Our findings demonstrate that TED can increase the survival rate of Vero cells infected with the HSV-1 virus, suggesting that TED has potential antiviral effects on HSV-1. Additionally, a plaque assay, which quantifies infectious viral particles by the number of cellular plaques that appear under dye staining after virus infection of cells, revealed that TED significantly suppressed HSV-1 replication, as evidenced by the reduced plaque formation rate in Vero cells. To further confirm the therapeutic efficacy of TED against HSV-1 *in vivo*, we established an animal model through intrastromal HSV-1 inoculation in mice. The HSK model mice exhibited characteristic disease manifestations, including progressive emaciation, circular gait patterns, elevated ocular surface clinical scores, marked corneal opacity, and extensive neovascularization. Histopathological analysis revealed significant corneal stromal degeneration and necrosis with substantial inflammatory cell infiltration. However, TED administration effectively attenuated these symptoms, indicating the notable protective effects of TED against HSV-1 infection in mice. To the best of our knowledge, this study represents the first investigation into the antiviral activity of a tectorigenin derivative (TED), demonstrating its great potential as a therapeutic agent for HSV-1-induced ocular diseases.

HSK represents an immunopathological disorder in which inflammatory cytokines play pivotal roles in driving disease progression and determining clinical severity. HSV-1 infection triggers the production of interferon (IFN), a cytokine that suppresses viral replication. IFN-α and IFN-γ, key members of the interferon family, are essential for controlling HSV-1 and antiviral defence in HSK^[28]^. However, their prolonged activation exacerbates stromal inflammation, leading to corneal opacification, thickening, and pathological angiogenesis^[29]^. In this study, we found that treatment with TED reduced the expression of IFN-α and IFN-γ in mouse corneas. In addition to IFNs, studies have demonstrated that the pharmacological regulation of inflammatory cytokines alleviates HSK progression. Here, TED administration exhibited dual immunomodulatory effects: suppression of inflammatory mediators (TNF-α, IL-1β) and enhancement of the anti-inflammatory cytokine IL-10. Collectively, these findings demonstrate that TED ameliorates HSK through dual mechanisms: by downregulating pathogenic IFN-α/IFN-γ responses and proinflammatory cytokine production while upregulating anti-inflammatory cytokine expression. This finding is consistent with the established therapeutic effects of tectorigenin in various inflammatory diseases, including liver fibrosis, fulminant hepatic failure, and *Dabie bandavirus*^[7,15, 30]^.

Like those in other inflammatory diseases, the secretion of IFN and inflammatory cytokines in HSK is closely associated with the TLR-mediated innate immune signalling pathway. To date, TLR-1, -2, -3, -4, -5, -7, -8, and -9 have been demonstrated to be expressed on corneal epithelial cells^[31]^. Among these TLRs, TLR2, TLR3, and TLR9 are extremely critical for HSK progression because they regulate IFN and inflammatory cytokine secretion^[32]^. Specifically, the HSV-1 glycoproteins gB/gH/gL activate TLR2-mediated pathways, whereas TLR3 and TLR9 sense virus-associated nucleic acids during infection, collectively exacerbating HSK immunopathology. In this study, HSV-1 infection significantly increased the expression of TLR2, TLR3, and TLR9 in mouse corneas, whereas TED treatment effectively reversed these alterations. Consistent with the changes in TLR2, TLR3, and TLR9 expression, the downstream adaptor proteins of these TLRs exhibited similar responses to both HSV-1 infection and TED treatment. Specifically, TED suppressed both the TRAF6-NF-κB cascade downstream of TLR2/9 and the IRF3-NF-κB axis downstream of TLR3. The suppression of TLR2/3/9 signalling by TED aligns with its inhibition of IFN-α/γ and proinflammatory cytokines (TNF-α, IL-1β) in this study. Collectively, these findings demonstrate that TED ameliorates HSK immunopathology through coordinated suppression of the TLR2/3/9 signalling network and its downstream effectors (TRAF6-NF-κB and IRF3-NF-κB), thereby attenuating both IFN-α/γ overproduction and excessive inflammatory cytokine responses.

While innate immunity provides the first line of defence against HSV-1 infection, the adaptive immune system plays an equally critical role in controlling HSK pathogenesis. Multiple immune cell populations coordinate these adaptive defences, with dendritic cells (DCs) serving as key bridges between innate and adaptive immunity^[33]^. Our results revealed marked infiltration of DCs in the cornea following HSV-1 infection, suggesting the triggering of virus-specific adaptive immune responses^[34]^. On the basis of the well-established role of DCs in activating T-cell-mediated immunity, we subsequently quantified CD4^+^ and CD8^+^ T-cell populations in the cornea given the central role of CD4^+^ T cells in orchestrating corneal immunopathology during HSK^[35]^, along with the dual antiviral functions of CD8^+^ T cells—which can eliminate HSV-1-infected cells by inducing apoptosis and suppress viral replication via IFN-γ secretion in response to infection^[36]^. Our results revealed no detectable CD4^+^ or CD8^+^ T-cell infiltration in normal corneas by immunofluorescence staining, whereas the number of infiltrating CD4+ or CD8+ T cells increased significantly after HSV-1 infection. These findings align with previous studies in which HSV-1 was inoculated into scarified BALB/c mouse corneas, followed by immunohistochemical detection of CD4^+^ and CD8^+^ T cells^[37]^. To trace the origin of these T-cell responses, we further examined the DLNs. Notably, HSV-1 infection induced a consistent downwards trend in both CD4⁺ and CD8⁺ T-cell numbers in DLNs, although only the reduction in CD4⁺ T cells reached statistical significance (*P* < 0.05). This drainage pattern strongly suggests active T-cell migration from DLNs to the infected cornea. To investigate whether TED could modulate this DC‒T-cell axis, we assessed its effects on both the corneal and DLN compartments. Strikingly, TED treatment (1) reduced DC infiltration in the cornea; (2) attenuated T-cell recruitment to the cornea; and (3) reversed HSV-1-induced T-cell depletion in DLNs, with CD4⁺ T-cell numbers recovering to 99% of control levels (*P* < 0.001). Most notably, TED treatment induced selective CD4⁺ T-cell recovery in DLNs, increasing the CD4⁺/CD8⁺ ratio to 2.61±0.08 compared with 2.07±0.23 in HSV-1-infected controls (*P* < 0.05), indicating preferential modulation of CD4⁺ T-cell homeostasis^[38]^. Since reduced corneal CD4⁺ T-cell infiltration is correlated with attenuated HSK severity, the preferential modulation of CD4⁺ T-cell homeostasis in DLNs by TED likely contributes to the therapeutic efficacy of TED by rebalancing local and systemic adaptive immunity.

In conclusion, this study demonstrated that TED, a tectorigenin derivative, exerts multifaceted therapeutic effects against HSV-1-induced HSK through coordinated modulation of innate and adaptive immunity. The key findings include the following: (1) Direct antiviral activity: TED suppresses HSV-1 replication *in vitro*, enhancing infected cell survival and reducing viral plaque formation. (2) Innate immune regulation: TED inhibits TLR2/3/9-mediated signalling, attenuating pathogenic IFN-α/γ and proinflammatory cytokine (TNF-α, IL-1β) production while increasing the anti-inflammatory cytokine IL-10. (3) Adaptive immune modulation: TED rebalances T-cell responses by (a) reducing DC-driven T-cell priming and corneal infiltration. (b) Preferential restoration of CD4⁺ T-cell homeostasis in DLNs. (c) Limiting corneal immunopathology through decreased CD4⁺ T-cell recruitment. These mechanisms collectively suppress the viral load and mitigate inflammation-induced corneal damage. Given its dual antiviral and immunomodulatory properties, TED represents a promising candidate for treating drug-resistant or recurrent HSK.

## Acknowledgements

We thank the Experimental Animal Centre of Sichuan Academy of Traditional Chinese Medicine and the Institute of Pharmacy of Sichuan Academy of Traditional Chinese Medicine for supporting this study.

## Transparency declarations

None to declare.

## Author contributions

J.ZHO. and L. ZHA. designed the study. K.N., M.Y., H.ZHA., H.M., and C.Y. performed the experiments. W.Y., Y.W., and H.X. participated in the data collection. K.N. and Y.H. analysed the data and wrote the manuscript.

## Data availability

The data that support the findings of this study are available from the corresponding author upon reasonable request.

